# Linking functional and structural brain organisation with behaviour in healthy adults

**DOI:** 10.1101/2024.07.04.602076

**Authors:** Natalie J Forde, Alberto Llera, Christian Beckmann

**Affiliations:** Radboud University Medical Centre, Donders Centre for Brain, Cognition and Behaviour, Nijmegen, Netherlands

**Keywords:** Linked ICA, healthy variation, brain-behaviour associations, multimodal, function-structure relationship, brain organisation

## Abstract

Multimodal data integration approaches, such as Linked Independent Component Analysis (LICA), increase sensitivity to brain-behaviour relationships and allow us to probe the relationship between modalities. Here we focus on inter-regional functional and structural organisation to determine if organisational patterns persist across modalities and if investigating multi-modality organisations provides increased sensitivity to brain-behaviour associations.

We utilised multimodal magnetic resonance imaging (MRI; T1w, resting-state functional [fMRI] and diffusion weighted [DWI]) and behavioural data from the Human Connectome Project (HCP, n=676; 51% female). Unimodal features were extracted to produce individual grey matter density maps, probabilistic tractography connectivity matrices and connectopic maps from the T1w, DWI and fMRI data, respectively. DWI and fMRI analyses were restricted to subcortical regions for computational reasons. LICA was then used to integrate features, generating 100 novel independent components. Associations between these components and demographic/behavioural (n=308) variables were examined.

15 components were significantly associated with various demographic/behavioural measures. 2 components were strongly related to various measures of intoxication, driven by DWI and resemble components previously identified. Another component was driven by striatal functional data and related to working memory. A small number of components showed shared variance between structure and function but none of these displayed any significant behavioural associations.

Our working memory findings provide support for the use of fMRI connectopic mapping in future research of working memory. Given the lack of behaviourally relevant shared variance between functional and structural organisation, as indexed here, we question the utility of integrating connectopic maps and tractography data.

## Introduction

Understanding the neural correlates of behaviour is challenging, both in healthy variation and pathological phenotypes. However this endeavour is vital as it provides the fundamental knowledge upon which better treatments are developed and clinical decisions based. Individual variation that often hampers classical approaches may be the key to unlocking our understanding of brain-behaviour relationships once appropriate approaches are implemented. One such approach is that of using various sources of information integrated together to give a fuller picture of underlying processes. The assumption with multimodal approaches is that underlying neurobiological variation (that relates to behaviour) influences multiple different indices of the brain as we measure it. Thus by integrating information from multiple sources we gain increased sensitivity to detect associations with behaviour.

In recent years linked independent component analysis (LICA), an extension to standard ICA that integrates multimodal data through a shared mixing matrix, has been utilised in neuroimaging studies to great effect. An initial paper by Groves et al (Groves et al. 2012) demonstrated its utility in linking multimodal brain phenotypes, based on structural measures, with demographic information such as age and sex. Additionally, former studies using LICA have been sensitive to concussion related brain phenotypes (Manning et al., n.d.), attention deficit/ hyperactivity disorder severity (Francx et al. 2016) and autistic diagnosis (Oblong et al. 2023). Furthermore Llera et al recently highlighted a positive-negative spectrum of behavioural measures associated with a multimodal phenotype, again based on structural measures (Llera et al. 2019). When functional connectivity was also included it added little in terms of variance explained. Although this showed little benefit to including functional data to the model, we propose that it may be worthwhile when different forms of information are extracted from the functional signal prior to integration with LICA.

For instance, it has long been known that the brain has behaviourally relevant topographic organisations and regions may have multiple overlapping functional organisations (Kaas 1997; Jbabdi, Sotiropoulos, and Behrens 2013). Recently, various methods have been developed to investigate the ‘gradual’ organisation of the brain, see (Watson and Andrews 2022) for an overview and comparison of methods. These methods are termed ‘connectopic mapping’. The method we implement here employs a non-linear manifold learning technique in a select region of interest (ROI) to extract one or more connectopic maps (Haak, Marquand, and Beckmann 2017). These represent the organisation of connectivity within the ROI based on the connectivity to the rest of the cortex. This method has proven useful in identifying behaviourally relevant organisations in primary motor and visual cortices (Haak, Marquand, and Beckmann 2017), subcortical structures (Marquand, Haak, and Beckmann 2017; Przeździk et al. 2019; Oldehinkel et al. 2022) and association cortices (Faber et al. 2020). These functional topographies allow for the multiplicity of brain regions as well as their continuous organisation. We propose that these functional topographies are more likely to relate to structural organisation than data aggregated over regions like in traditional functional connectivity analyses. Additionally, we extend this focus on regional organisation to our diffusion weighted Imaging (DWI) data analysis. Previous studies using LICA have utilised DWI data processed through restricted voxel-wise based methods (i.e. tract-based spatial statistics [TBSS] (S. M. Smith et al. 2006)) and tensor models of water diffusion (Llera et al. 2019; Francx et al. 2016). Jeurrissen and colleagues have previously shown that the tensor model is inadequate in the majority of white matter voxels due to the complexity of the white matter architecture and the low resolution of imaging (Jeurissen et al. 2013). We therefore use a non-tensor based model and probabilistic tractography approach to model white matter organisation. Moreover we examine the fine grained pattern of how a region is structurally connected to the rest of the brain rather than aggregating a signal across the whole region of interest (ROI).

We hypothesise that these advanced approaches to unimodal data extraction coupled with multimodal integration via LICA will provide us with additional sensitivity to detect brain-behaviour relationships akin to and in addition to those already described (Llera et al. 2019). We recently successfully implemented a similar protocol in the analysis of a large sample of autistic and non-autistic individuals where we focussed on autistic traits and diagnosis (Oblong et al. 2023). Here we take a broader behavioural perspective in a large neurotypical sample to expand our understanding of brain-behaviour relationships. Moreover our focus on regional organisation of brain regions could help shed more light on the structure-function relationship.

## Methods

### Data

We utilised the large high quality Human Connectome Project (HCP) healthy young adult data (Glasser et al. 2013) (S1200 release). Only subjects with all of the required modalities were included (T1-weighted, rsfMRI (only reconstruction type ‘r227’ and DWI). Subjects were also excluded if head coil instabilities were noted (n=73). This resulted in a sample of N = 682. A further 6 subjects were excluded due to processing failures. Resulting in a final sample of 676 subjects (51% female). See supplementary table 1 for demographic information.

### Unimodal feature extraction

See figure 1 for an overview of the unimodal feature extraction and multimodal integration. T1-weighted MRI data were processed with the Computational Anatomy Toolbox (CAT 12) from SPM, an extension to the voxel based morphometry (VBM) pipeline (J. Ashburner and Friston 2000; John Ashburner and Friston 2005). This pipeline includes improved denoising and brain extraction procedures. All individual T1 images were affine aligned before being segmented, normalised and bias-field-corrected generating individual segmented maps (grey matter, white matter and CSF). Diffeomorphic registration of all images to a standard grey matter template was then achieved with DARTEL (John Ashburner 2007). Finally, a FWHM Gaussian smoothing kernel of 9.4 mm was applied (sigma = 4 mm).

**Figure 1.**
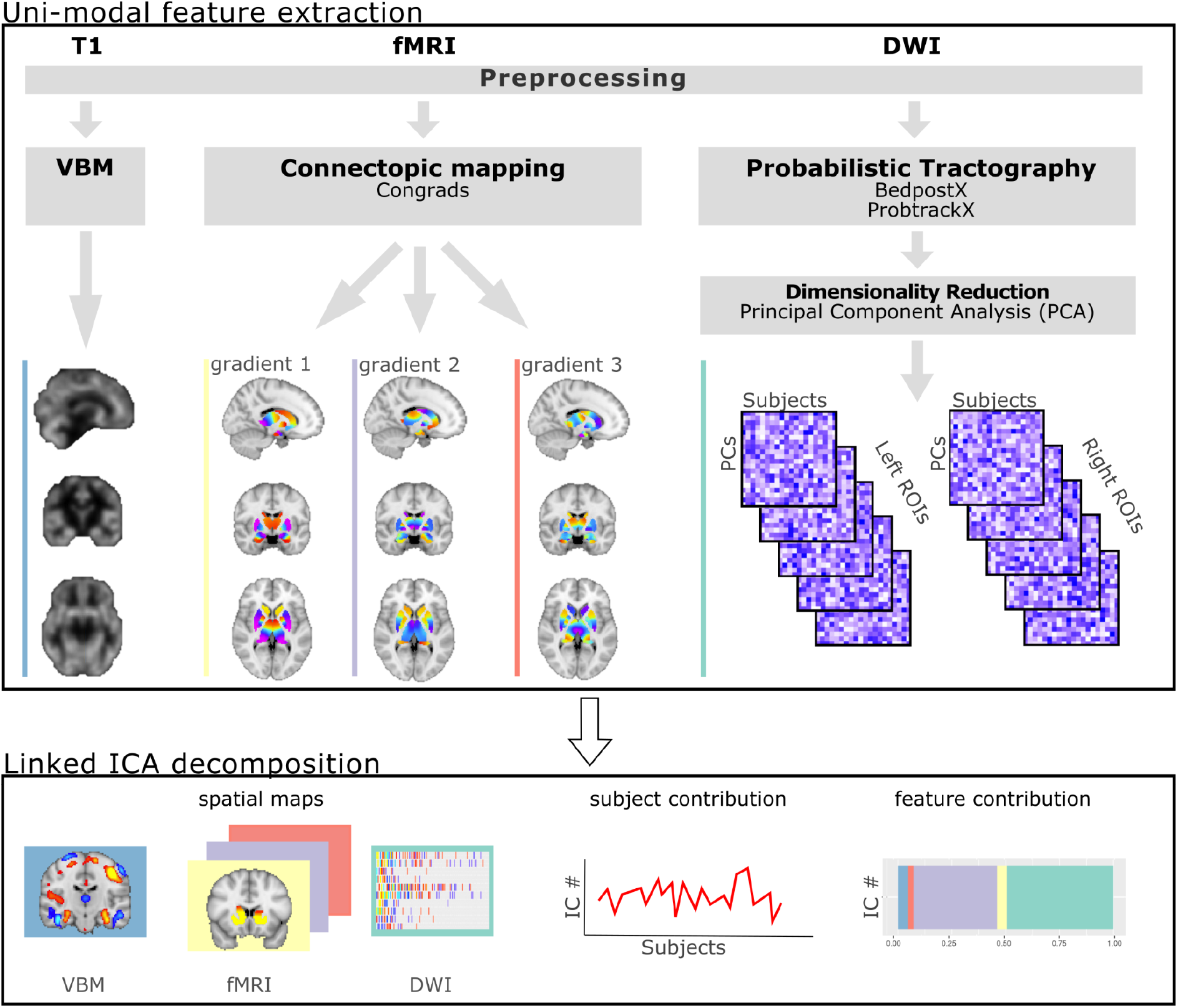
Schematic overview of the processing pipeline for unimodal feature extraction and integration with linked independent component analysis (LICA). fMRI - functional MRI, DWI - diffusion weighted imaging, T1 - T1-weighted MRI, VBM - Voxel based morphology, PCs - Principal components, IC - independent component.

### ROI selection for DWI and RS analysis

Due to computational restrictions analysis had to be limited to select ROIs. Based on the previous literature utilising connectopic mapping showing behavioural associations with subcortical structures (Marquand, Haak, and Beckmann 2017; Przeździk et al. 2019; Oldehinkel et al. 2022) we focus here on subcortical ROIs. ROIs were isolated from the Harvard-Oxford atlas as provided by FSL. The striatum ROI was formed by combining the Nucleus Accumbens, Putamen and Caudate structures per hemisphere.

### Diffusion weighted imaging (DWI)

DWI data were acquired in a minimally preprocessed form from HCP, which included correction for movement and eddy current induced artefacts (Glasser et al. 2013). Data were then processed with Bayesian Estimation of Diffusion Parameters Obtained using Sampling Techniques (BEDPOSTX) and probabilistic tractography (ProbtrackX) (Behrens et al. 2007). ProbtrackX was performed with seeding from select ROIs and a list of target atlas ROIs to generate a voxel by target ROI connectivity matrix for each seed ROI. The target list was composed of cortical regions from the multi-modal parcellation developed on the HCP data (Glasser et al. 2016) and subcortical ROIs from the Harvard-Oxford atlas. A MNI 2×2×2 mm^3^ template was used for atlas regions to ensure spatial correspondence across subjects. Principal component analysis (PCA) was used on voxel by atlas matrices to reduce the dimensionality of the data. Matrix rank-1 PCs were kept for analysis in LICA. Participant-by-PC matrices for the different seed ROIs were stacked to produce one DWI input for LICA.

### Resting state functional MRI (rs-fMRI)

Rs-fMRI data were also acquired in a minimally preprocessed form directly from HCP, this included preprocessing with ICA-FIX to identify and remove components related to motion and imaging artefacts (Glasser et al. 2013). The HCP protocol included the collection of 4 rs-fMRI scans, 2 each in opposing phase encoding directions. Here we selected ‘REST1’ RL and LR per subject to balance encoding direction and maximise the number of subjects included. Additionally 2 different reconstruction algorithms were used by HCP for the rsfMRI data, for consistency we chose to use only one of these, r227.

We smoothed the data with a FWHM Gaussian kernel of 5.9 mm (sigma 2.5 mm). We then performed connectopic mapping to generate topographic maps of each ROI determined by their functional connectivity to the rest of the cortex (Haak, Marquand, and Beckmann 2017). Reference gradients were produced by averaging the gradients from 20 randomly selected subjects. All subject gradients were then checked against these to ensure consistent ordering and direction (i.e. not flipped). Gradients 1 to 3 were selected for further analysis. Gradients (1, 2 or 3) from the different ROIs were combined to give three inputs for LICA, one for each gradient. Research has shown evidence for 2 overlapping organisations in certain brain regions (Jbabdi, Sotiropoulos, and Behrens 2013; Haak, Marquand, and Beckmann 2017). The choice to focus on the first 3 gradients instead of 2 was based on recent research showing higher order modes may also be of interest (Oldehinkel et al. 2022).

### Multimodal integration

LICA is an extension to ICA that allows for the integration of multi-modal data linked through a shared mixing matrix (Groves et al. 2011). For each independent component (IC) isolated the algorithm provides a set of spatial maps (one per modality), a vector of the loading weights showing the contribution of each modality to that component and finally a vector describing the contribution of each subject. This final vector is the subject loadings per IC which can be used to investigate the relationship between the brain phenotypes and demographic/behavioural measures. We generated 100 components for this analysis. We calculated a multimodal index (MMI) per IC as previously described (Francx et al. 2016).

### Demographic & behavioural data

A wide array of demographic and behavioural data were utilised. From all data that was available if more than 5% of data were missing then the measure was excluded. Otherwise missing data were median imputed. Additionally, variables were excluded where variance was extremely low such that >95% of subjects had the same value. Finally, race and ethnicity variables were excluded. This left 308 demographic and behavioural measures. For a full list of measures investigated see supplement. Data included were in line with the previous study from Llera et al (Llera et al. 2019).

### Statistical Analysis

Permutation Analysis of Linear Models (PALM (Winkler et al. 2014)) was used to test the correlations between demographic/behavioural measures and subject loadings from linked-ICA components while accounting for non-independence of subjects due to relatedness with exchangeability blocks. Additional data manipulation (e.g. multiple comparison correction) and graph generation were performed in R (V3.5.1, (R Core Team 2013)). Multiple comparison correction was implemented with the false discovery rate (FDR) procedure (Benjamini and Hochberg 1995).

### Additional analyses

Data integration and analyses were repeated, as above, with subsets of the modalities; structural only (VBM and DWI) or functional (Gradients 1-3). Additionally, as the connectopic maps were originally scaled (0-1) we repeated analyses without scaling the maps, see supplement for more details and outcomes.

## Results

Figure S1 shows the feature contributions to each component and the associated MMI. There was one component reflecting a strong balanced combination of the input modalities, IC 92 (MMI > 0.8). About a fourth of the ICs provided a high level of multimodality showing MMI greater than 0.5. The modalities that tended to show the greatest shared variance within ICs were VBM and DWI or the functional gradients with each other. However, a few ICs showed shared variance between the functional and structural modalities, see supplement for further details. None of these highly multimodal function-structure ICs displayed significant behavioural associations.

### Behavioural associations

15 components were significantly associated with demographic/behavioural measures (Figure S7 summarises these findings). Here we highlight 3 of interest (Figure 2). IC’s 9 and 15 showed significant associations with multiple demographic/behavioural measures including many related to intoxication. Additionally, IC14 was related to multiple measures of working memory.

**Figure 2.**
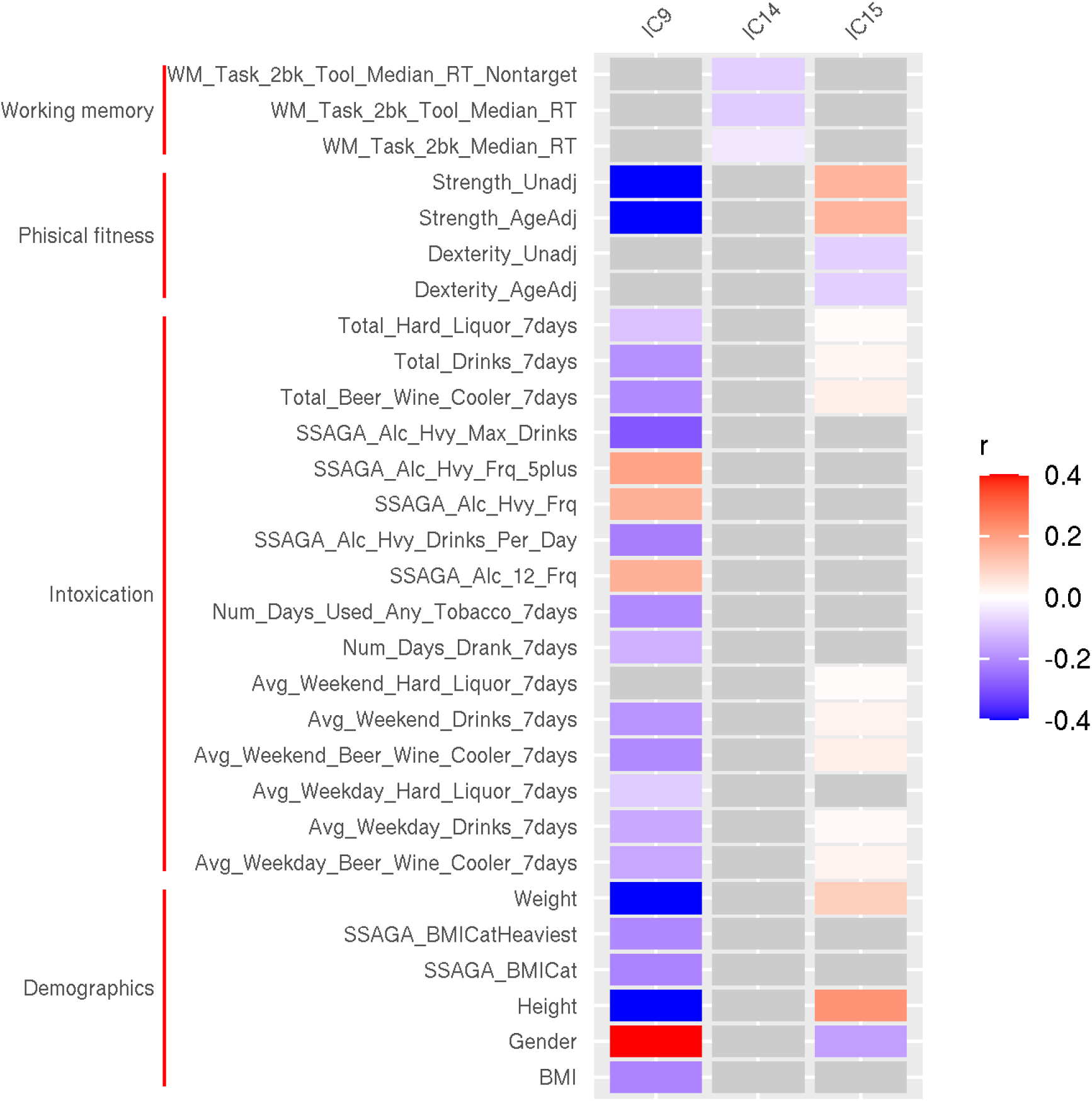
significant behavioural associations with ICs 9, 14 and 15 Significant behavioural associations are coloured according to their correlation coefficient (r).

IC9 and IC15 were dominated by DWI data (97% and 96%, respectively) with additional small contributions from VBM (3%, 4%, respectively; Figure 3). All ROIs from the DWI data appear to contribute mostly with their first few PCs, suggesting a very global connectivity association. IC14 on the other hand was dominated by fMRI gradient 3 (97%; Figure 3). Upon visual inspection we see that the effect is strongly localised to the left striatum. No other feature contributed more than 1%. Full results and spatial contributions to each of the other significant ICs are presented in the supplementary material.

**Figure 3.**
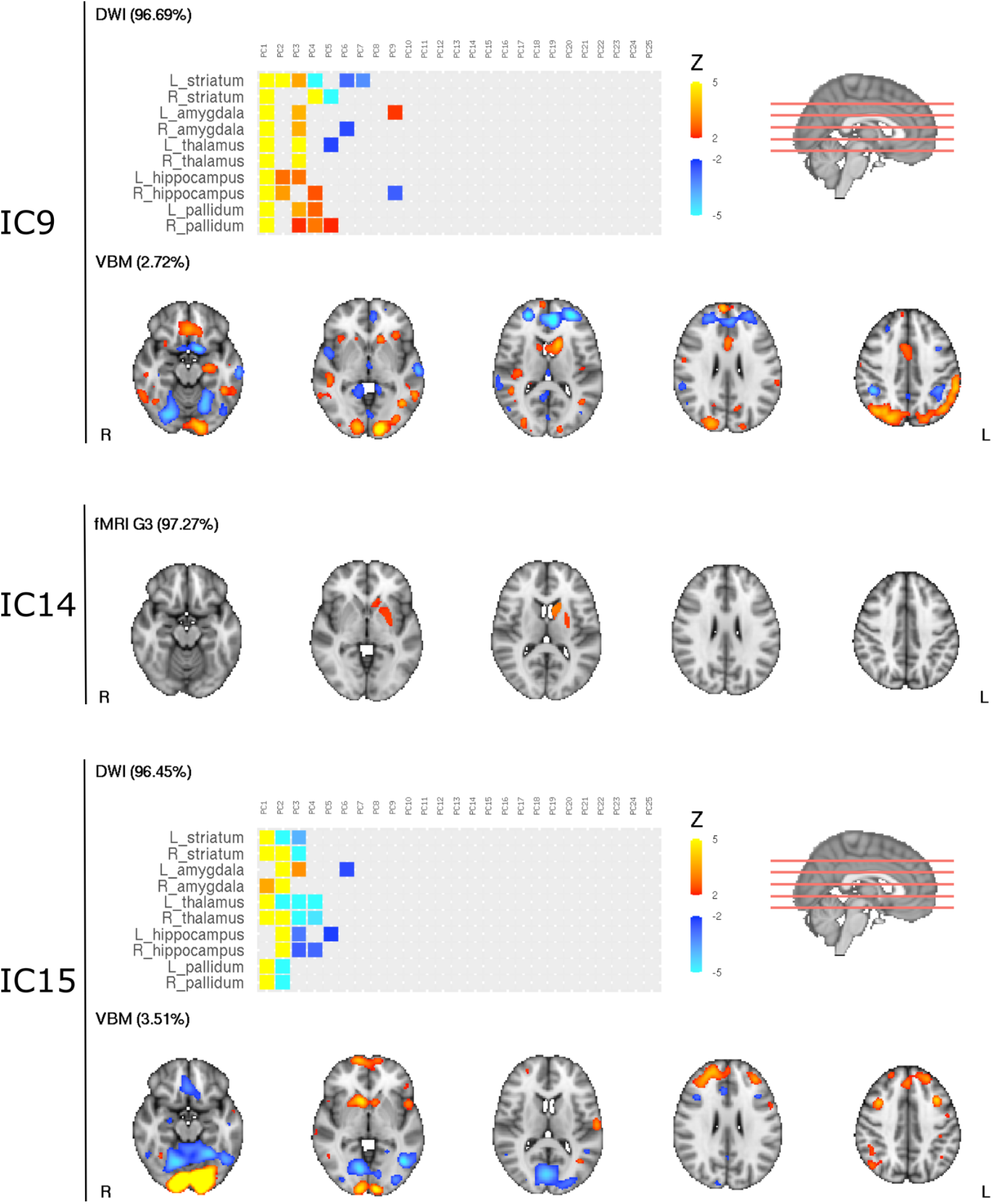
spatial maps for ICs 9, 14 and 15. PC’s 1-25 of the DWI data are visualised. |z| > 2 is shown. Only features contributing more than 2% are shown.

### Sensitivity/Validation

To ensure our LICA model wasn’t failing to pick up more shared variance between structure and function due to the scaled nature of the connectopic maps we reran LICA with unscaled connectopic maps. This had little influence on the distribution of structure-function feature variance in ICs. Additionally, LICA was also run with structural and functional data separately to determine if separately they explain behavioural variance better than together. These analyses provided negligibly different ICs to those in the main analysis. Details of these additional analyses are presented in the supplementary material.

## Discussion

This study has integrated MRI data from three MR modalities including structural and functional data. Advanced unimodal processing of both functional and diffusion MR data provided the opportunity to interrogate the potential structure-function relationship between modalities when modelled in a framework that allows for continuous patterns of organisation within ROIs, along with behavioural associations of these brain phenotypes. Numerous ICs generated were strongly linked to demographic and behavioural measures, however, the majority of these showed little, if any, shared variance between structural and functional modalities. A small number of ICs generated do capture a high degree of shared structural and functional variance, however, none of these structure-function ICs were significantly related to any of the behavioural metrics investigated.

A wide array of papers have been published which investigate the structure-function relationship (Shen et al. 2015; C. J. Honey et al. 2009; Christopher J. Honey et al. 2007; Fields 2008) or structure-function coupling in the brain (Gu et al. 2021; Baum et al. 2020). These vary greatly in approach and aims but none, that we are aware of, take into account the functional multiplicity of regions and their (potentially) gradual organisation. Here we attempted to address this by modelling the functional signal as a continuum within each ROI and including multiple such gradients from each ROI. These gradients have previously been shown to relate to different aspects of behaviour (Marquand, Haak, and Beckmann 2017; Przeździk et al. 2019) and recently dopaminergic transporter availability (Oldehinkel et al. 2022). Additionally, most former analyses of dMRI in structure-function studies have been limited to signal aggregated to ROIs, thus, ignoring any potential variation in organisation within regions. Our analysis utilised connectivity patterns of voxels within subcortical ROIs to the rest of the brain. Allowing intra-regional organisations, that may relate to functional organisations, to be captured. Despite these advances we did not isolate brain phenotypes, composed of functional and structural signals, that relate to behavioural measures. However, a few ICs generated showed a high degree of shared variance between functional and structural modalities. These may nonetheless be of interest for understanding the structure-function relationship itself or potentially relate to behavioural measures not indexed in the current study (see supplement for more in depth review of those ICs). This is inline with our previous study in autism using similar methods where we saw few ICs in the decomposition that showed shared variance between structural and functional modalities (Oblong et al. 2023).

There were however numerous ICs (15) related to at least one demographic or behavioural measure. The majority of these associations were with demographics such as sex, height, weight and strength. This is in agreement with previous findings (Llera et al. 2019; Groves et al. 2012). However, we found only 1 component related to age, little in comparison to previous papers, but this is likely related to our small age range in contrast to the Groves paper and our exclusion of morphological modalities where age effects were mostly previously localised (i.e. pial area).

Two components from the current analysis correspond closely with components isolated previously by Llera et al (see Table S3, Figure S12-S14) (Llera et al. 2019). All these components have multiple behavioural associations, most notably with measures of intoxication. The components from the current study show fewer behavioural associations which may relate to the small number of ROIs that were utilised for the DWI analysis rather than the whole brain approach in the former study. Despite the focal nature of the DWI modality analysis both of our ICs were overwhelmingly driven by the DWI data with minimal contribution from VBM (our structural only decomposition reproduces these components very closely - see supplement for details). This is in contrast to the study by Llera where the loading of VBM was much stronger in both ICs (55% and 97%, respectively). This could imply that our method of DWI analysis, despite not being whole-brain, is a sensitive marker of brain variation that relates to behaviour. Moreover, extending analysis to more regions (or whole brain) may further increase the sensitivity to brain-behaviour associations. In contrast to the similarity in findings from these ICs, it is noteworthy that we do not replicate Llera et al’s primary finding of interest - that of the positive-negative spectrum of behavioural associations with one structural component. This positive-negative behavioural mode was initially proposed by Smith and colleagues (Stephen M. Smith et al. 2015) and related to functional connectivity. While later studies demonstrated the spatial topography or spatial overlap of functional networks was driving the behavioural association more than functional coupling (Bijsterbosch et al. 2018,2019). Finally, Llera et al replicated the positive-negative mode using structural only data and showed functional data had little impact in explaining behavioural variability beyond that explained by structural variability (Llera et al. 2019). Here we compared our results to those of Llera et al whose study closely matched the current design. One of our components (IC81) was highly correlated (rho = -0.47, *p*_*perm*_<0.001, Figure S15B), based on an overlap of n=207 participants, with the previously reported component (IC6). Interestingly, in our structural only decomposition, our IC5 relates to Llera’s IC6 (rho =-0.45, *p*_*perm*_<0.001, Figure S15C). However, we see no significant behavioural associations with our components. Uncorrected associations bear some similarity to those found previously and are presented in the supplement (Figure S15 A and D) for comparison with the former study. Again our lack of significant findings may relate to the focal nature of our DWI analysis or potentially to our exclusion of some modalities that formerly contributed - Llera’s component was contributed to by pial area (4%), jacobian difference (7%) and cortical thickness (7%) as well as the DWI metrics (60% total) and VBM (22%). Further details of the comparison can be found in the supplementary material.

Another component presented here (IC14), was related to three measures of working memory and almost entirely dominated by the contribution from the subcortical gradient 3. Which in turn was dominated by the left striatum. Striatal gradient 3 has, to our knowledge, not been previously investigated in relation to behavioural measures. However, the role of the striatum in working memory is well known (Emch, von Bastian, and Koch 2019; Wager and Smith 2003). Interestingly, striatal dopamine functioning has been linked to working memory capacity (Landau et al. 2009) and task related functional connectivity appears to mediate this association (Nour et al. 2019). Moreover, recent work from our colleagues has shown a striatal gradient to reflect the dopaminergic transporter availability seen with DaT SPECT imaging (Oldehinkel et al. 2022). I must caution that the gradients in these papers are not identical as the ROIs used to define them were slightly different. Nonetheless, our finding does suggest that the novel use of resting state fMRI gradients for the mapping of dopaminergic projections may be useful in the investigation of intra-individual variability in working memory, as well as in ageing and Parkinson’s Disease as already proposed.

Our study was limited by computational restraints that resulted in the inclusion of only subcortical ROIs. Given current findings and previous literature we expect that the inclusion of additional regions in future research would produce ICs with more significant brain-behaviour associations. Despite our efforts to incorporate unimodal functional and diffusion modelling of our data to best capture potential intra-regional organisations these may nonetheless have been limited - either by not including a sufficient number of functional gradients to capture relevant signal in the functional analysis or the coarseness of the atlas used for target regions in the DWI analysis which still aggregates signal to some degree. Additionally, our broad exploratory approach utilising all available behavioural data required an enormous amount of multiple comparison correction potentially limiting our sensitivity to detect behavioural associations in comparison to taking a more refined *a priori* approach and restricting analysis to select behavioural measures.

In summary, we have performed a novel integration of multimodal MRI data utilising regional brain organisations for the first time. Unfortunately, we failed to see significant correspondence between functional and structural brain organisation but do identify multiple brain phenotypes related to demographic and behavioural measures. Among these are components related to intoxication and working memory. Additionally, we provide support for the use of fMRI connectopic mapping in future research of working memory.

## Acknowledgements

Data were provided by the Human Connectome Project, WU-Minn Consortium (Principal Investigators: David Van Essen and Kamil Ugurbil; 1U54MH091657) funded by the 16 NIH Institutes and Centers that support the NIH Blueprint for Neuroscience Research; and by the McDonnell Center for Systems Neuroscience at Washington University.

