## Supplemental Material for "Linking functional and structural brain organisation with behaviour in healthy adults"

#### Table of Contents

**Supplementary methods** **3**
Table S1 Basic demographics 3
Additional analyses 3

**Supplementary results to main analysis** **3**
Figure S1 Modality contributions 4
Structure-function multimodal ICs 4
Figure S2 Modality contributions for select multimodal ICs 5
Figure S3 spatial maps for IC68 6
Figure S4 spatial maps for IC80 7
Figure S5 spatial maps for IC92 8
Figure S6 spatial maps for IC98 9
Behavioural associations 10
Figure S7 behavioural associations (all modalities) 10

**Supplementary results to additional analyses** **11**
Structural only analyses 11
Figure S8 behavioural associations structural only analysis 12
Table S2 correlations between all modality and structural components 12
Figure S9 behavioural associations; all modalities vs structural 13
Figure S10 spatial maps for IC1 and IC2 (structural) 14
Function only analyses 14
Figure S11 behavioural associations; all modalities vs functional 15
Non-scaled gradient analyses 15
Comparison with Llera et al (Llera et al., 2019) 15
Table S3 comparison with Llera et al; IC1 and IC2 16
Figure S12 subject courses from Llera et al; IC1 and IC2 vs current study 17
Figure S13 spatial maps from Llera et al; IC1 and IC2 18
Figure S14 spatial maps from IC9 and IC15 19
Table S4 comparison with Llera et al; IC6 20
Figure S15 behavioural associations and subject courses from Llera et al; IC6 21
Figure S16 spatial maps from Llera et al; IC6 22
Figure S17 spatial maps from IC81 (all modalities) and IC5 (structural) 23

References **24**

### Supplementary methods

**Table S1 Basic demographics**

| | Male | Female |
|---|---|---|
| n | 333 | 343 |
| Age, mean (sd) | 28 (3.7) | 29 (3.7) |
| FluidCog, mean (sd) | 106 (18) | 107 (17) |

##### Additional analyses

Data integration and analyses were repeated with subsets of the modalities; structural only (VBM and DWI) or functional.

Additionally, as the connectopic maps were originally scaled (0-1) we repeated analyses without scaling the maps. To determine the orientation and order of gradients we used the initial scaled gradients referenced against the 20 subject average. Re-ordering, where necessary, was then applied to the unscaled gradients. 'Flipping' of gradients, when necessary, was achieved by inverting the gradient (multiplying by -1).

### Supplementary results to main analysis

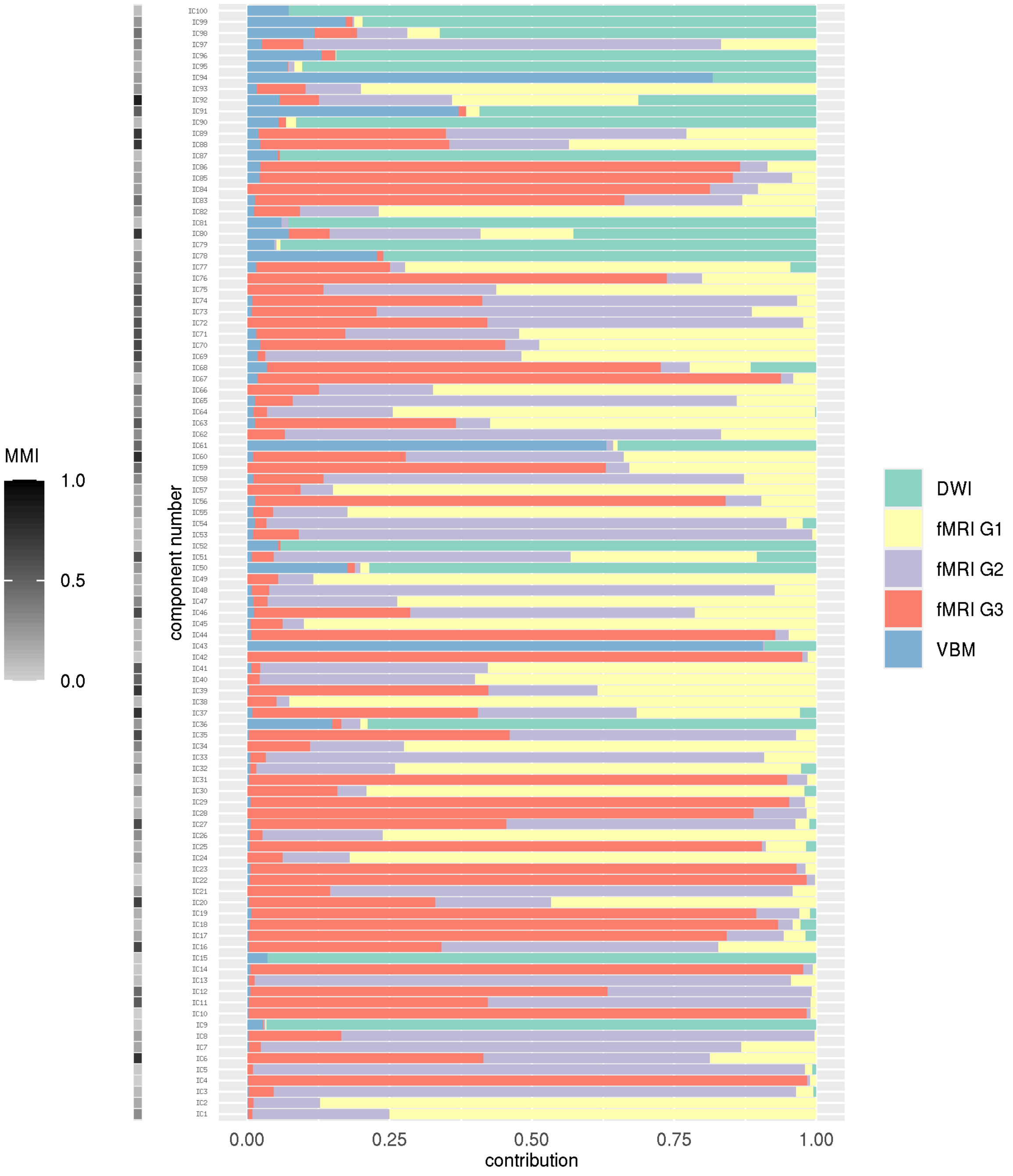


**Figure S1 Modality contributions**
Figure shows the modality contribution of all 100 ICs generated. MMI - multimodal index.

##### Structure-function multimodal ICs

The majority of multimodal components consisted of variance shared between either structural modalities (VBM and DWI) or functional gradients. However, a small number of ICs showed a greater degree of shared variance between structural and functional modalities. The modality contributions for these select ICs are shown in Figure S2. While figures S3-S6 show the spatial contributions for the same select ICs. Spatial convergence across modalities of contributing variance can be seen to some degree in all 4 ICs displayed. However, it is worth mentioning that DWI and fMRI data were restricted to subcortical ROIs so only those regions could contribute while the VBM data was whole brain.

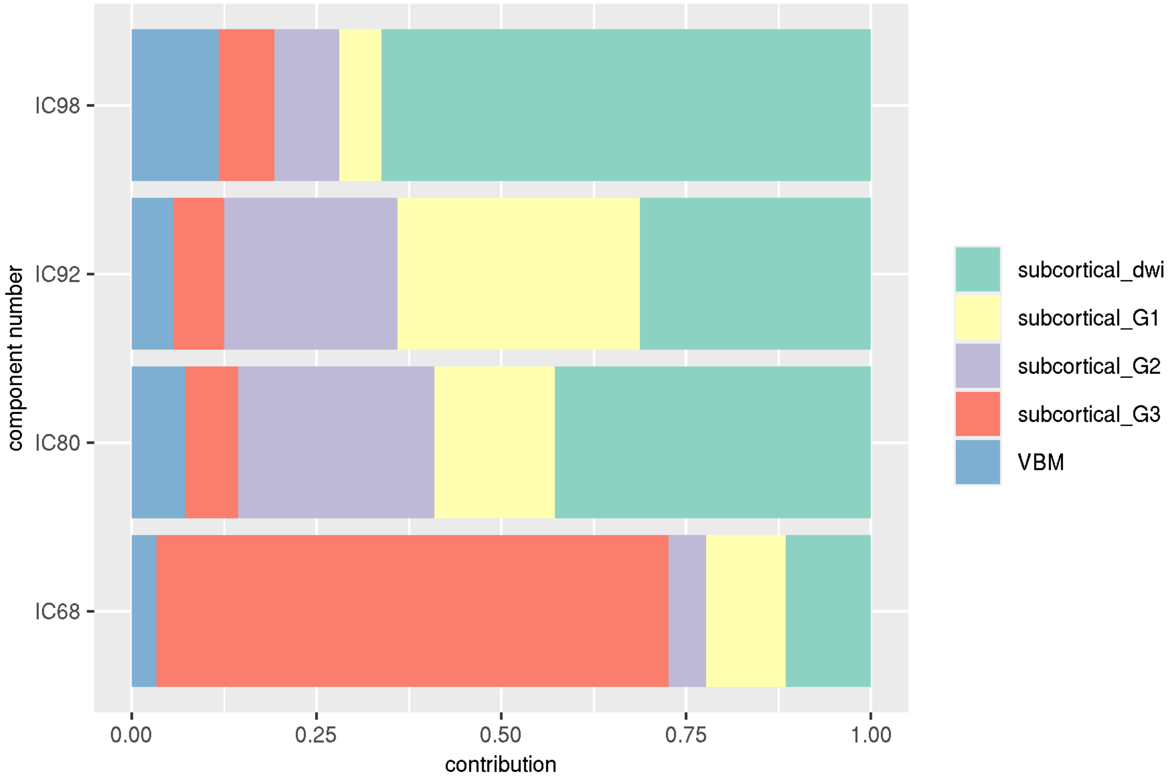


**Figure S2 Modality contributions for select multimodal ICs**
Figure shows the modality contribution of select ICs that display a high degree of shared variance between functional and structural modalities.

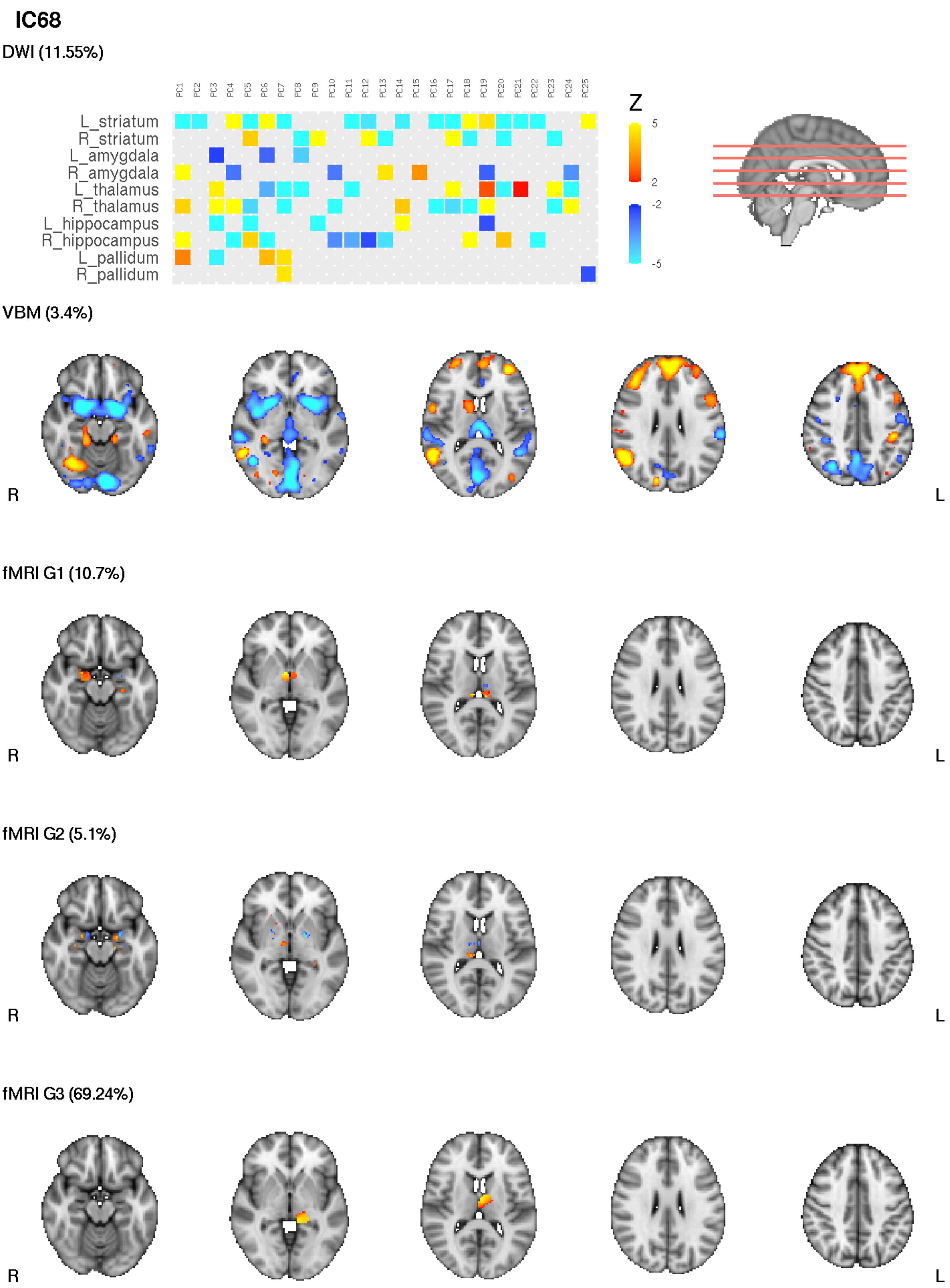


**Figure S3 spatial maps for IC68**

PC's 1-50 of the DWI data is visualised. |z| > 2 is shown.

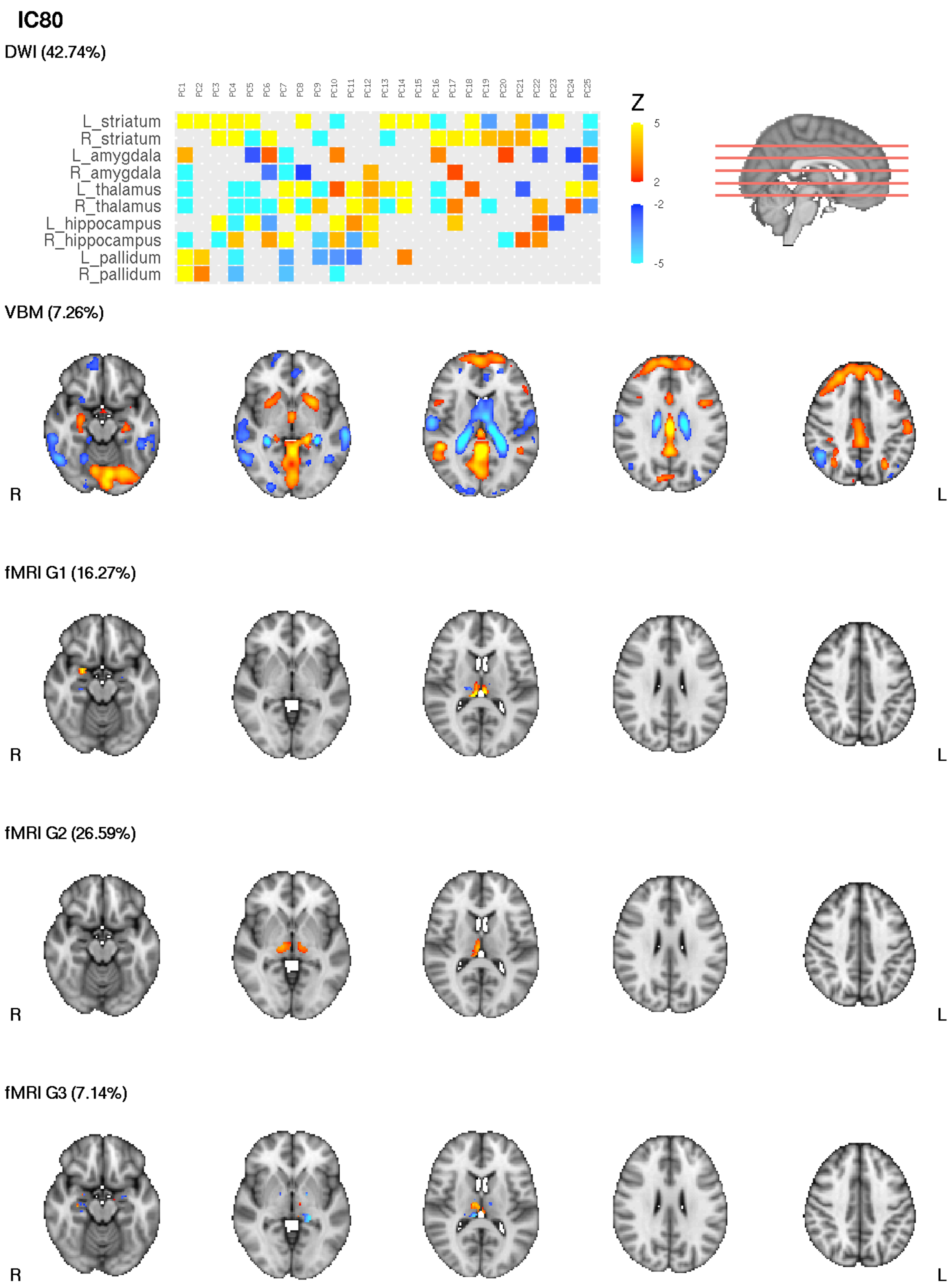


**Figure S4 spatial maps for IC80**
PC's 1-50 of the DWI data is visualised. |z| > 2 is shown.

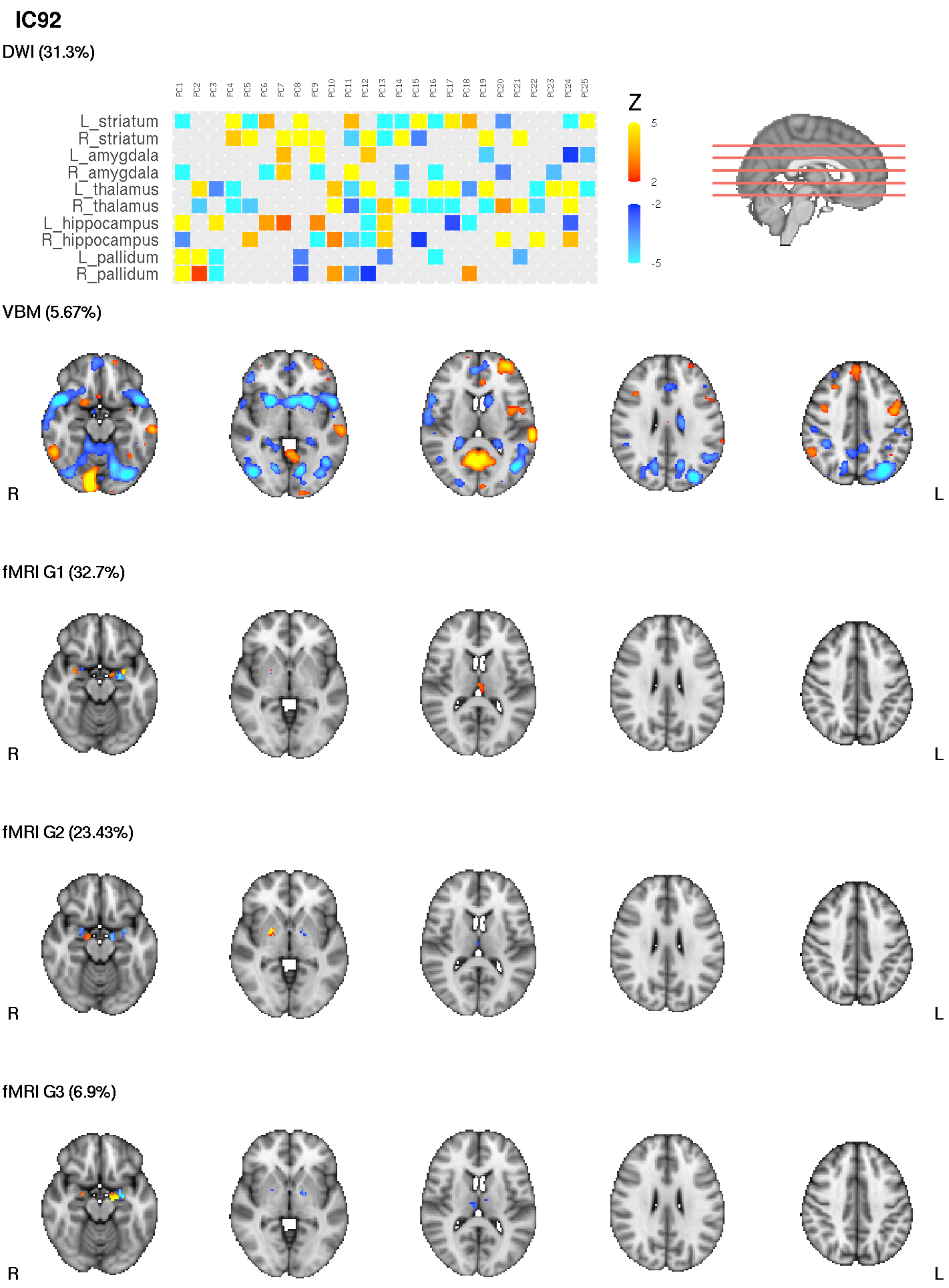


**Figure S5 spatial maps for IC92**

PC's 1-50 of the DWI data is visualised. |z| > 2 is shown.

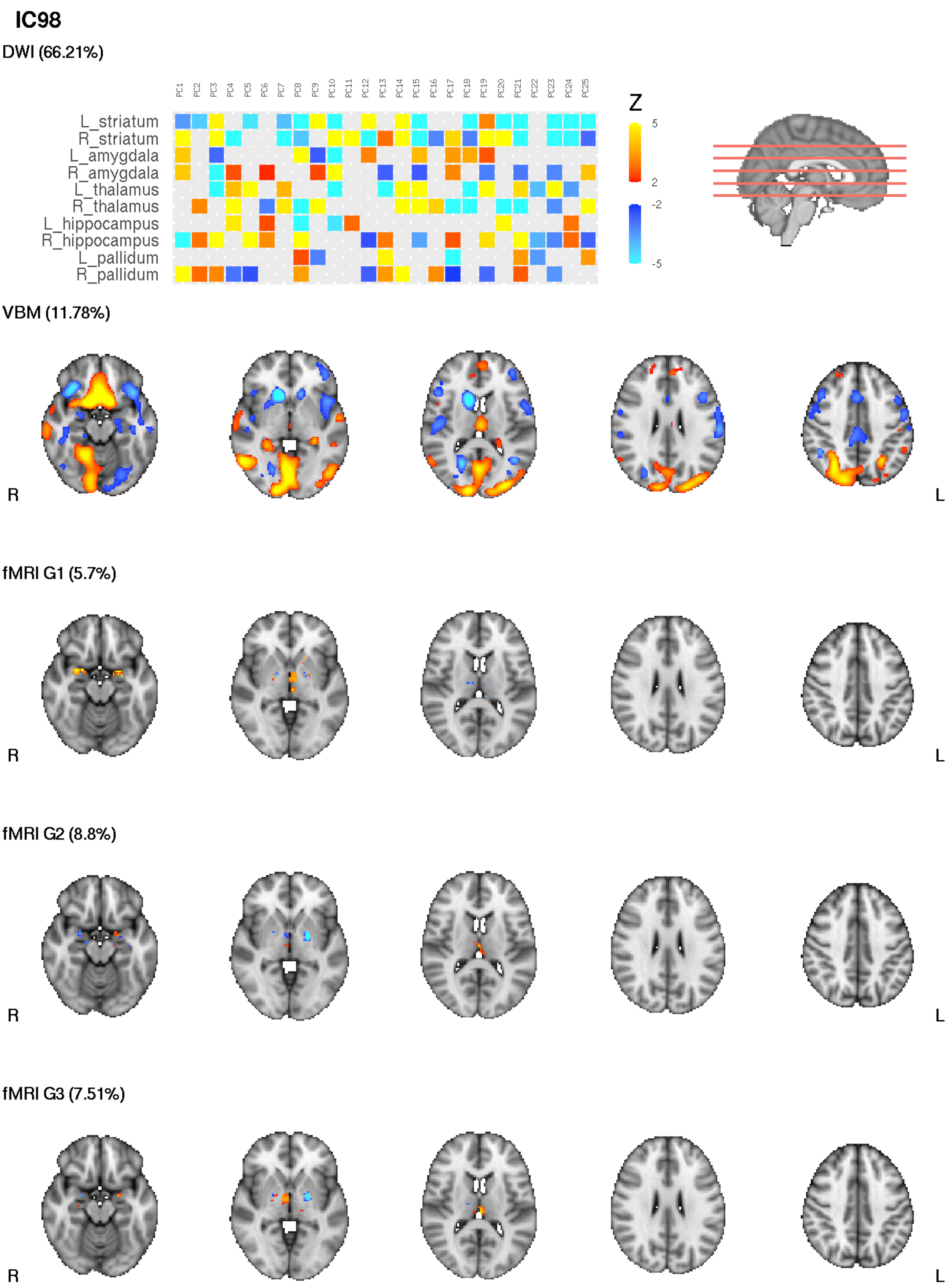


**Figure S6 spatial maps for IC98**

PC's 1-50 of the DWI data is visualised. |z| > 2 is shown.

#### Behavioural associations

In addition to ICs 9, 14 and 15 which we report in the main text, multiple other ICs were significantly associated with behavioural or demographic measures. All behavioural associations are summarised in Figure S7.

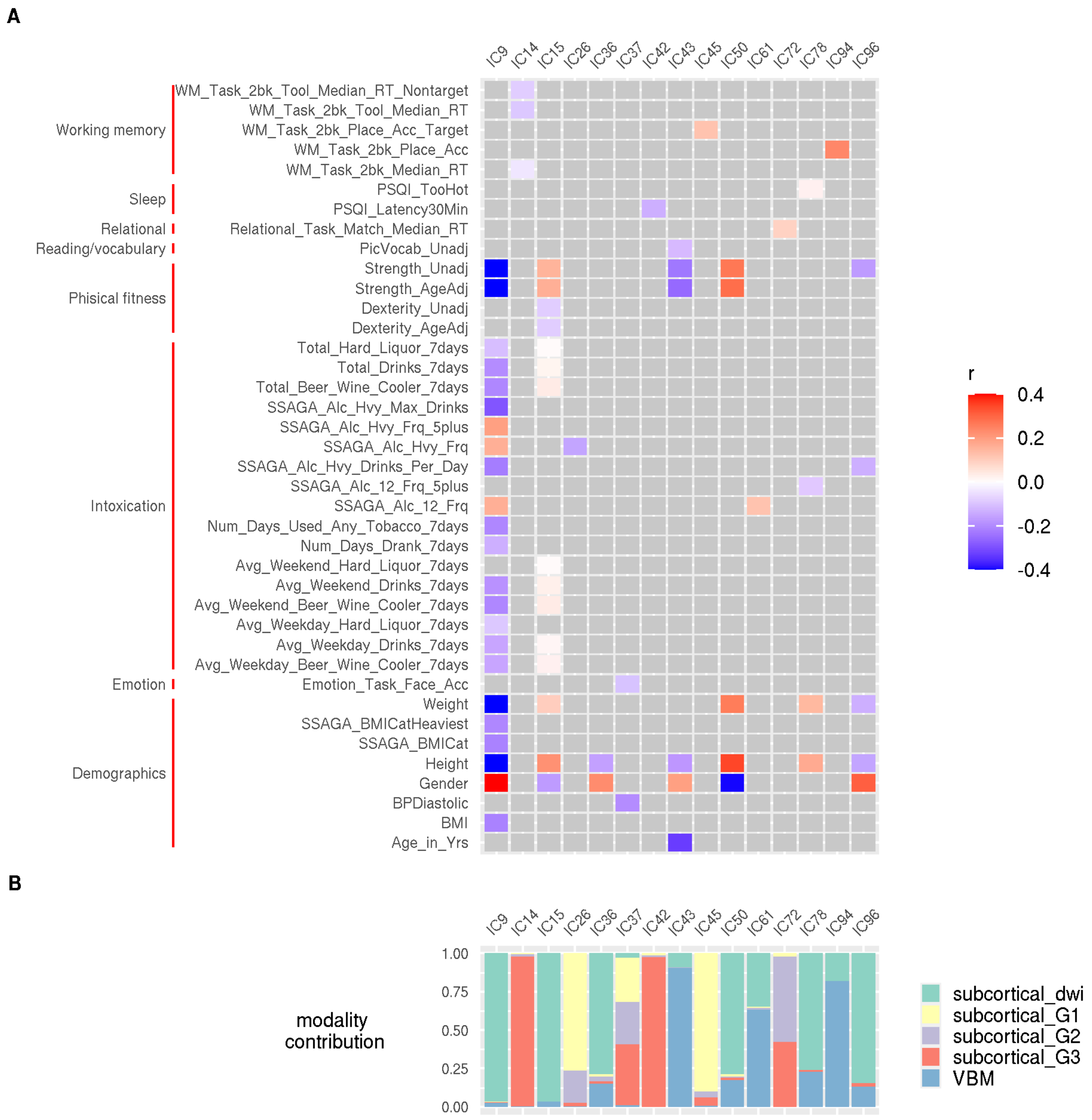


**Figure S7 behavioural associations (all modalities)**
(A) Significant behavioural associations from main analysis of all modalities are coloured according to their correlation coefficient (r). (B) shows corresponding modality contributions for these independent components (ICs).

### Supplementary results to additional analyses

#### Structural only analyses

Structural only analyses revealed an array of behavioural and demographic associations. These are summarised in Figure S8. Two structural components (IC1 and IC2) correlated strongly with IC9 and IC15 from the main analysis (rho = 0.99 and rho = 0.92, respectively, permutation p-values <0.001). These structural ICs also showed similar behavioural associations to the ICs from the main analysis (Figure S9). Additionally, the spatial patterns of these structural ICs correlated highly with those of IC9 and IC15 from the initial analysis (Table S2, Figure S10).

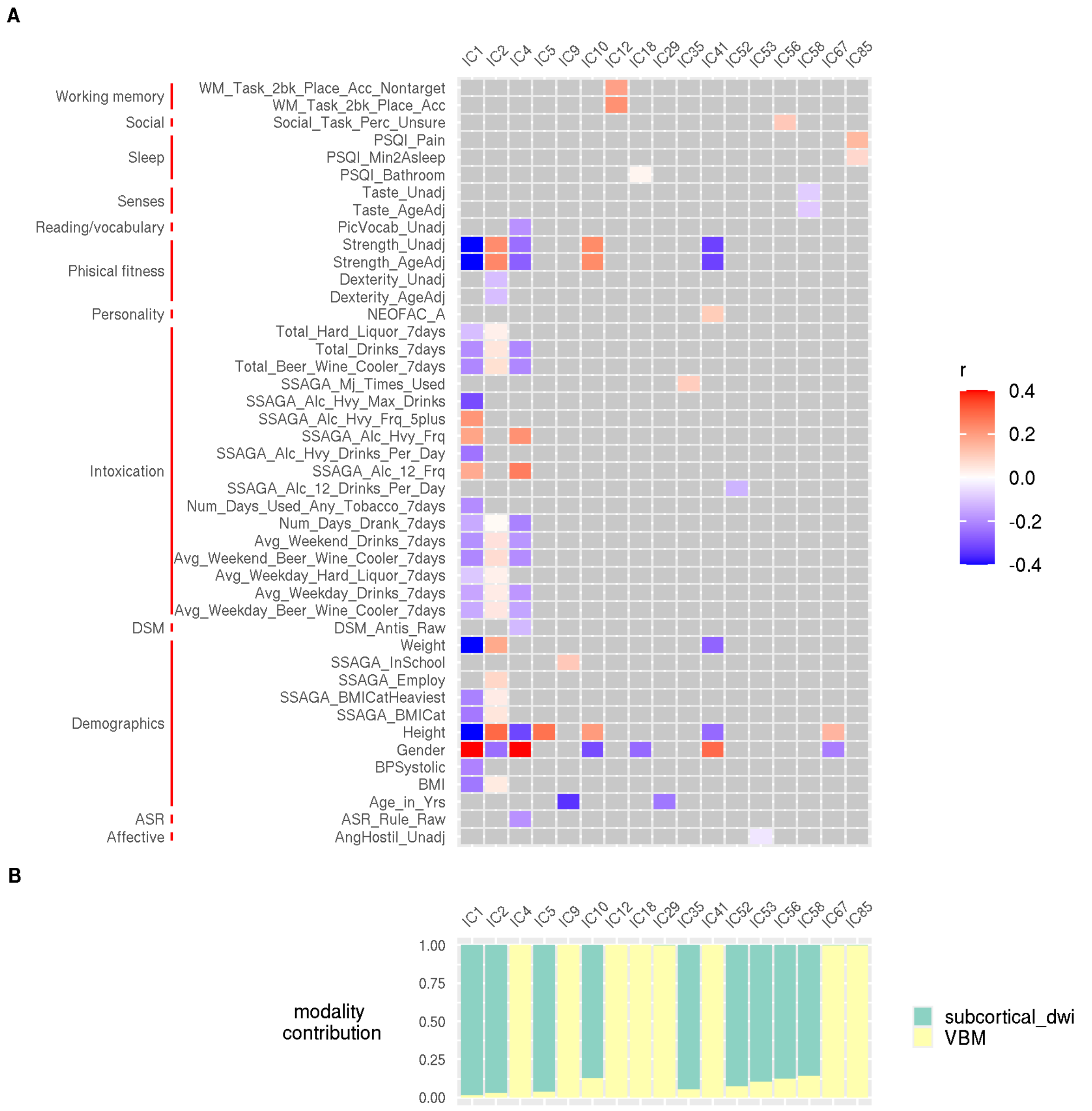


**Figure S8 behavioural associations structural only analysis**
(A) Significant behavioural associations from structural only analysis are coloured according to their correlation coefficient (r). (B) shows corresponding modality contributions for these independent components (ICs).

**Table S2 correlations between all modality and structural components**

| | Feature contributions | Subject course | DWI spatial maps | VBM spatial maps |
|---|---|---|---|---|
| All modalities IC9 | 97% DWI; 3% VBM | rho = 0.99, $p_{perm}$<0.001 | r = 0.89 | r = 0.69 |
| Structural IC1 | 99% DWI; 1% VBM | | | |

| | | | | |
|---|---|---|---|---|
| All modalities IC15 | 96% DWI; 4% VBM | rho = 0.92, $p_{perm}$ <0.001 | r = 0.75 | r = 0.89 |
| Structural IC2 | 97% DWI; 3% VBM | | | |

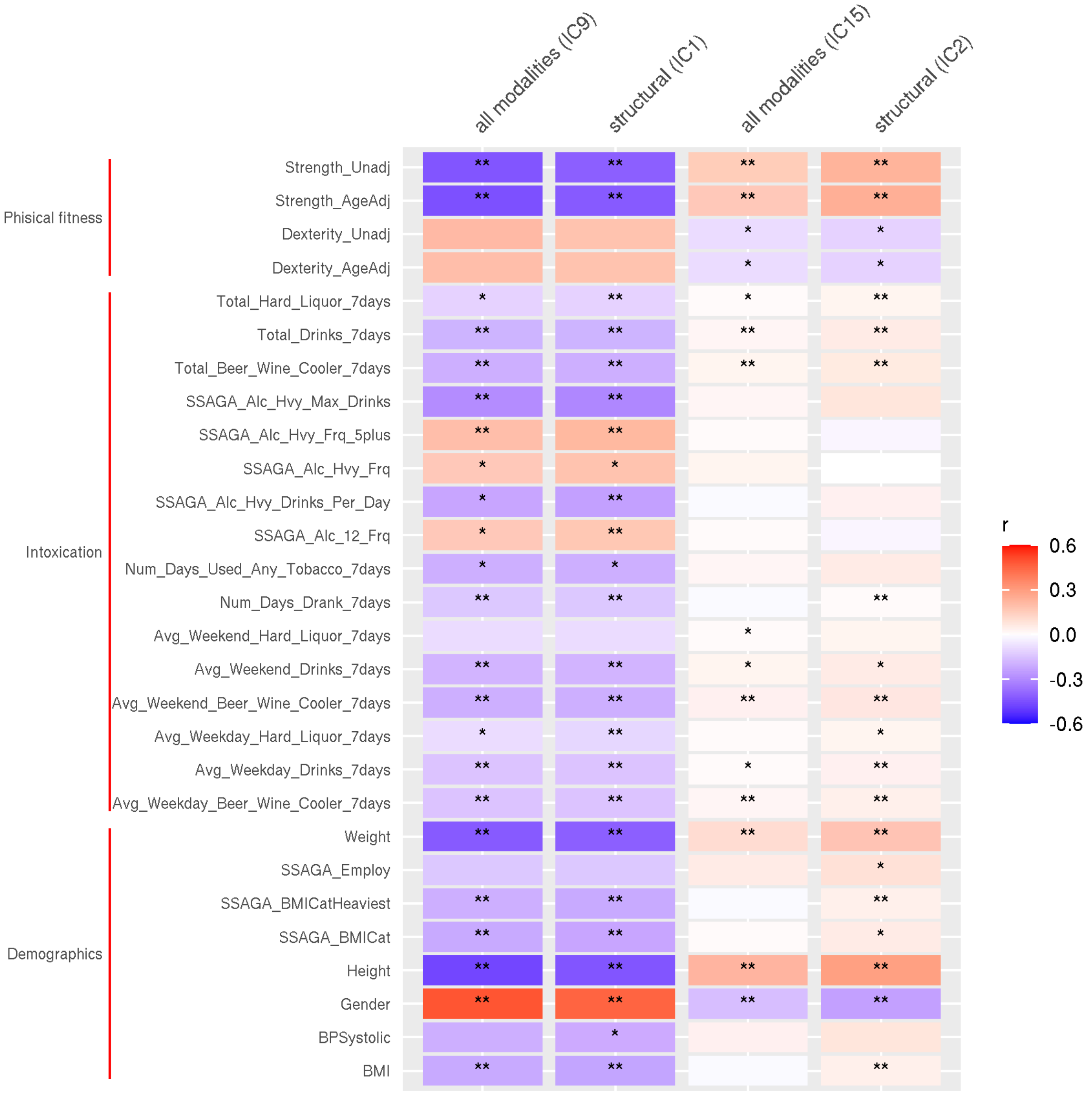


**Figure S9 behavioural associations; all modalities vs structural**

Select ICs from main and structural analyses that show similar behavioural associations. Tiles are coloured according to their correlation coefficient (r). * $p < 0.05$, ** $p < 0.01$, *** $p < 0.001$.

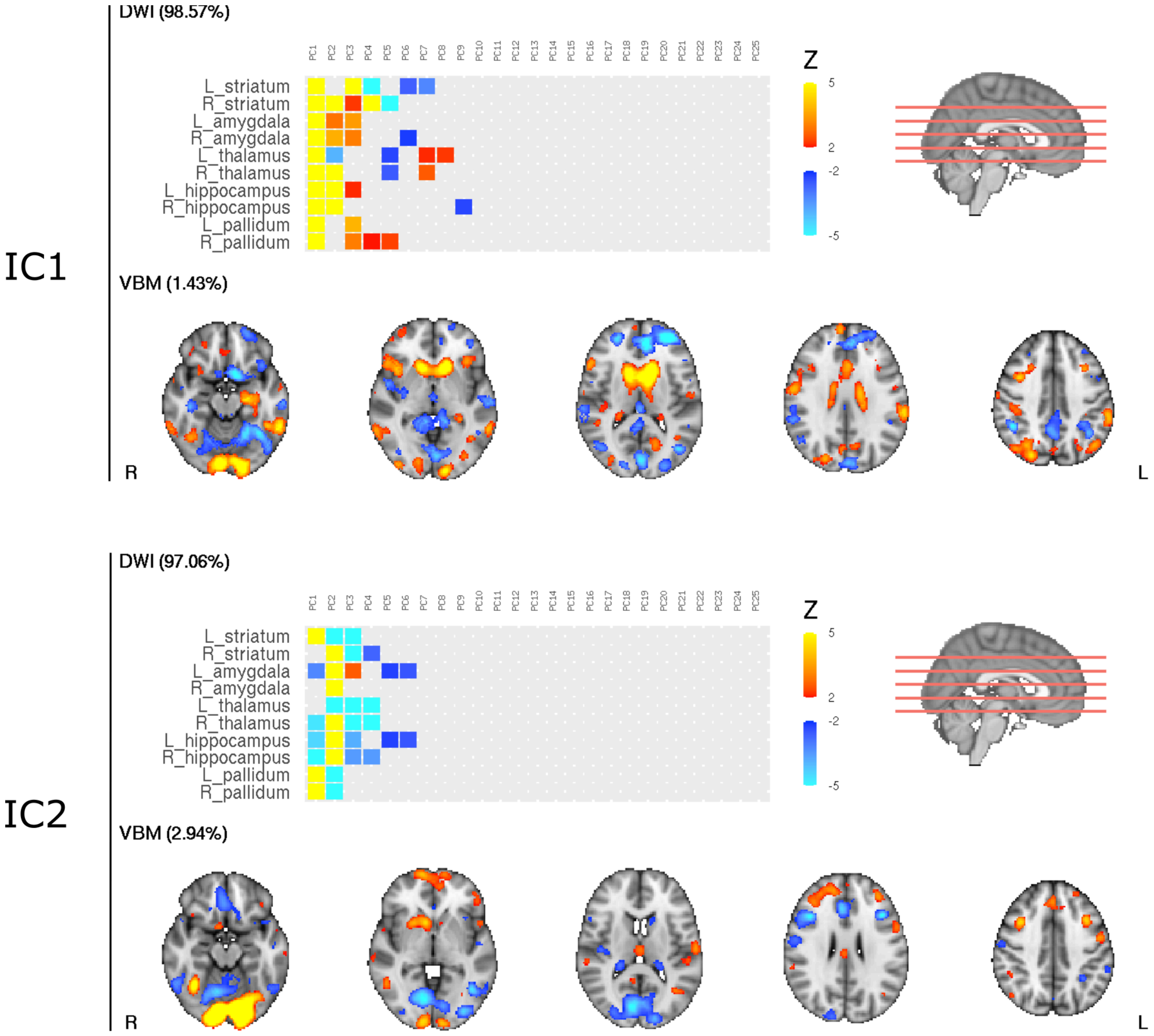


**Figure S10 spatial maps for IC1 and IC2 (structural)**
PC's 1-25 of the DWI data are visualised. |z| > 2 is shown. Only features contributing more than 2% are shown.

#### Function only analyses

Functional only data didn't reveal any significant behavioural associations when analysed on their own. One component (IC15) was strongly correlated with IC14 from the main analysis (subject courses: rho = 0.92, permutation *p* <0.001). Although no behavioural associations survived multiple comparison correction before correction similar behavioural associations to the main analysis were visible (Figure S11).

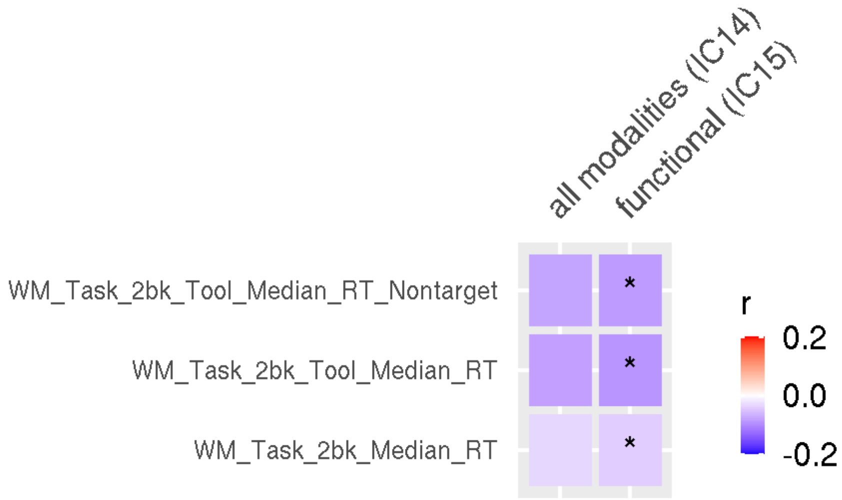


**Figure S11 behavioural associations; all modalities vs functional**
Select ICs from main and functional analyses that show similar behavioural associations. Tiles are coloured according to their correlation coefficient (r). * $p_{adj} < 0.05$

#### Non-scaled gradient analyses

Using non-scaled gradient data had little impact on findings, either in terms of shared variance across modalities or behavioural associations.

#### Comparison with Llera et al (Llera et al., 2019)

Components 1 and 2 from Llera et al correspond closely to components 9 and 15 from the current analysis (see Table S3 and Figure S12). Spatial maps (VBM and DWI only) from these components are displayed in Figures S13-14).

**Table S3 comparison with Llera et al; IC1 and IC2**

| | **All modalities IC9**<br>97% DWI; 3% VBM | **All modalities IC15**<br>96% DWI; 4% VBM |
|---|---|---|
| **Llera IC1**<br>10% DWI total (5% FA; 3% MD; 3% MO); 55% VBM; 2% JD; 2% CT; 30% PA | Subject courses<br>rho = -0.50, $p_{perm} < 1x10^{-16}$<br>VBM spatial maps<br>r = 0.14 | Subject courses<br>rho = 0.55, $p_{perm} < 1x10^{-16}$<br>VBM spatial maps<br>r = 0.09 |
| **Llera IC2**<br>0% DWI total (0% FA; 0% MD; 0 % MO); 97% VBM; 1% JD; 2% CT; 0% PA | Subject courses<br>rho = 0.36, $p_{perm} < 1x10^{-7}$<br>VBM spatial maps<br>r = 0.10 | Subject courses<br>rho = -0.31, $p_{perm} < 1x10^{-6}$<br>VBM spatial maps<br>r = 0.08 |

Percentages represent the modality contributions to the Independent Components (ICs). Correlations (Spearman) are calculated based on n=207 overlapping subjects from the two studies. Significance was tested by means of permutation (10000 permutations). Correspondence in the VBM spatial maps was calculated using FSL's fscc command. DWI - diffusion weighted imaging, VBM - voxel based morphometry, JD - Jacobian deformation, FA - fractional anisotropy, MD - mean diffusivity, MO - anisotropy mode, CT - cortical thickness, PA - pial area.

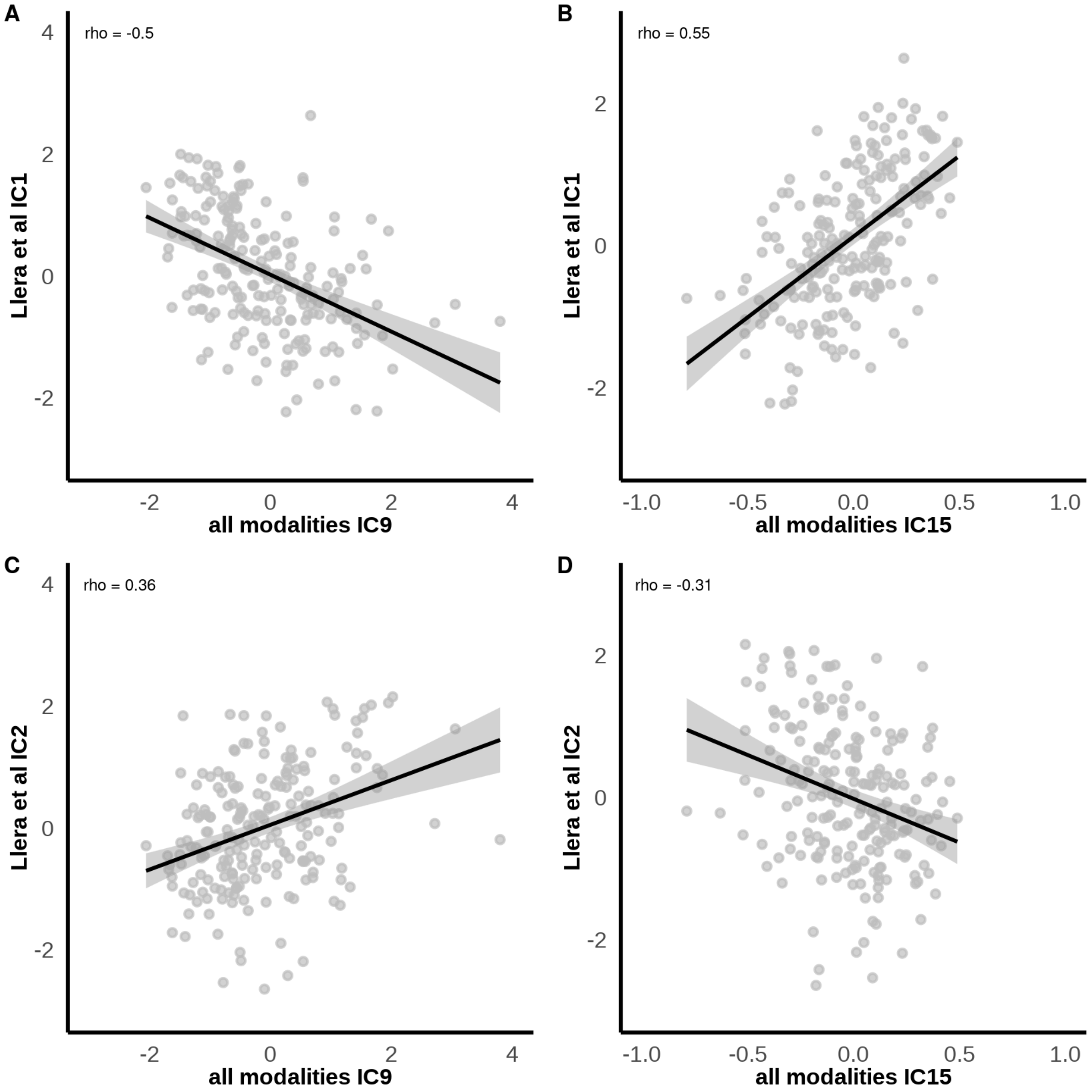


**Figure S12 subject courses from Llera et al; IC1 and IC2 vs current study**

All correlations (Spearman) were significant based on permutation testing ($p_{perm}$ <0.001). Calculations are based on n=207 individuals that overlapped between study cohorts. IC1 and IC2 are those reported by (Llera et al., 2019) and are seen to correspond closely to IC9 and IC15 from the current study (main analysis, all modalities included).

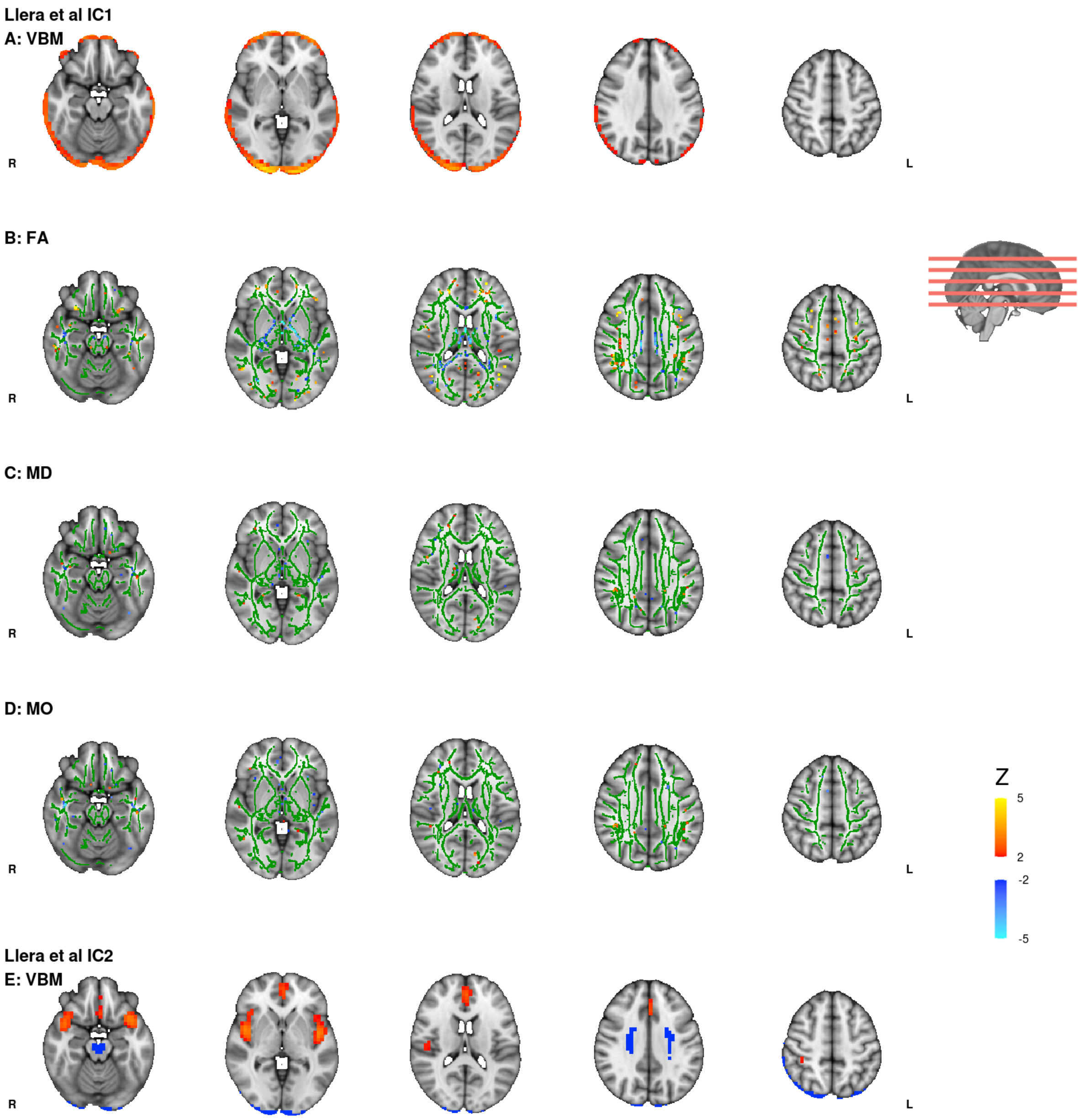


**Figure S13 spatial maps from Llera et al; IC1 and IC2**
Spatial maps, for select modalities, generated in the analysis by (Llera et al., 2019) are presented for IC1 and IC2. These are seen to correspond closely to IC9 and IC15 from the current study (main analysis, all modalities included). (A-D) relate to the VBM, FA, MD and MO modalities, respectively for IC1. (E) represents the VBM modality from IC2. Only VBM and DWI modalities are visualised and those contributing <3% were additionally excluded from visualisation. Colour gradient (blue-yellow) represents the Z score of the spatial map. Maps were thresholded at |z|>2. Sagittal brain slice shows location of axial slices used for each modality. Spatial maps are projected on the MNI T1 anatomical brain image for reference. DWI maps (FA, MD, MO) are displayed on top of the binarized FMRIB58 FA skeleton (green) as well as the MNI T1 template.

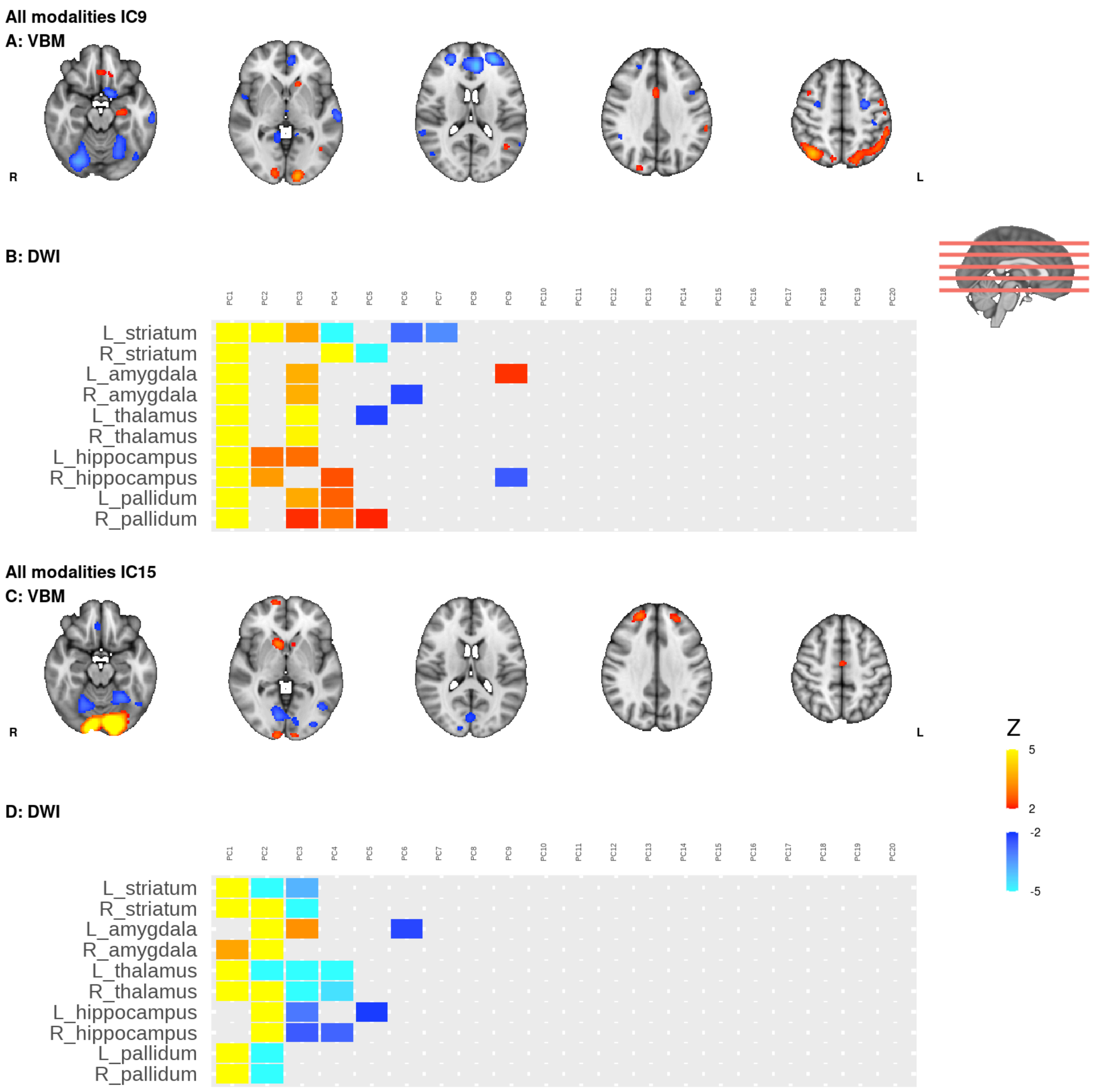


**Figure S14 spatial maps from IC9 and IC15**

Spatial maps, for select modalities, generated in the current study are presented for IC9 and IC15. These are seen to correspond closely to IC1 and IC2 from Llera et al. (A-B) relate to the VBM and DWI modalities, respectively for IC9. (C-D) similarly relate to the VBM and DWI modalities, respectively for IC15. Only VBM and DWI modalities are visualised. Colour gradient (blue-yellow) represents the Z score of the spatial map. Maps were thresholded at $|z|>2$. Sagittal brain slice shows location of axial slices used for VBM visualisation. VBM maps are projected on the MNI T1 anatomical brain image for reference. DWI maps only display PCs 1-20.

Component 6 from Llera et al, which was the component associated with a positive negative behavioural spectrum, was not reflected in our behaviourally significant components. The component from our analysis that correlates the most with IC6 in terms of subject contributions is IC81 (rho = -0.47, $p_{perm}$<0.001, Figure S15B). Additionally IC5 from the structural only analysis is significantly related to Llera's IC6 (rho = -0.45, $p_{perm}$<0.001, Figure S15C). Uncorrected behavioural associations for both these components are presented below for the purposes of comparison with the former study (Figure S15 A and D).

We find it interesting that the variance captured by these ICs differ so greatly in their respective position in their decompositions (6 and 5 when only structural data are included vs 81 when functional connectopic maps are also included). This suggests that the decomposition in the current main analysis, which includes connectopic maps and probabilistic tractography data, involves a high degree of variance that the decomposition utilises (n=80 components) before attributing the variance in question to a component. That is in stark contrast to the structural only decompositions reported in the Llera analysis and here which utilised only 5 or 4 components, respectively, before attributing this related variance to a component. Moreover, modality contributions (see table S4) reveal the greater weight of tractography based DWI preprocessing over tensor based maps in the decompositions.

Spatial maps (VBM and DWI only) from these components are displayed in Figures S16-17.

**Table S4 comparison with Llera et al; IC6**

| | **All modalities IC81**<br>93% DWI; 6% VBM | **Structural IC5**<br>96% DWI; 4% VBM |
|---|---|---|
| **Llera IC6**<br>58% DWI total (15% FA; 23% MD; 20% MO); 22% VBM; 6% JD; 7% CT; 4% PA | Subject courses<br>rho = -0.47, $p_{perm} < 1x10^{-16}$<br>VBM spatial maps<br>r = 0.08 | Subject courses<br>rho = -0.45, $p_{perm} < 1x10^{-16}$<br>VBM spatial maps<br>r = 0.14 |

Percentages represent the modality contributions to the Independent Components (ICs). Correlations (Spearman) are calculated based on n=207 overlapping subjects from the two studies. Significance was tested by means of permutation (10000 permutations). Correspondence in the VBM spatial maps was calculated using FSL's fscc command. DWI - diffusion weighted imaging, VBM - voxel based morphometry, JD - Jacobian deformation, FA - fractional anisotropy, MD - mean diffusivity, MO - anisotropy mode, CT - cortical thickness, PA - pial area.

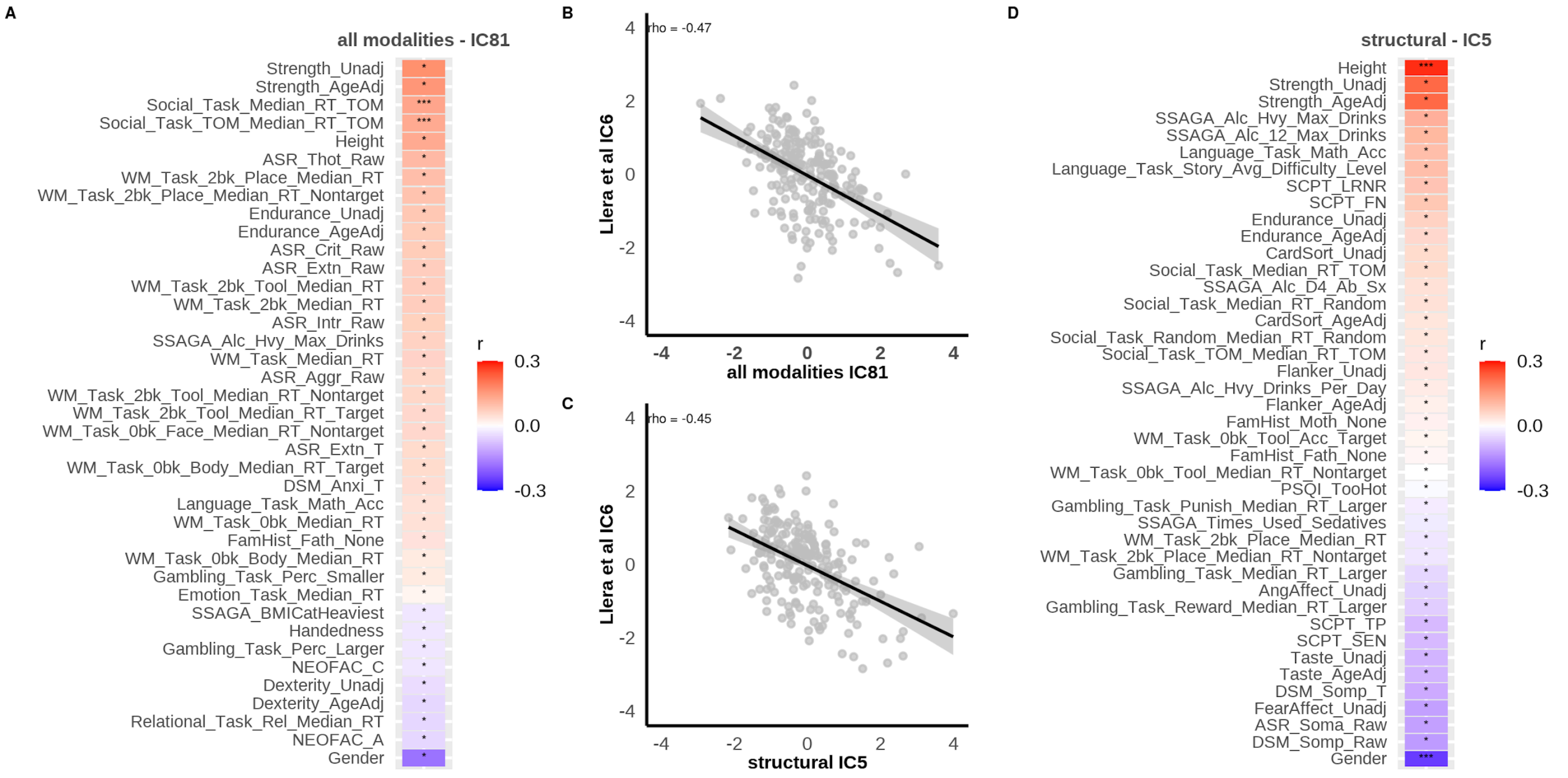


**Figure S15 behavioural associations and subject courses from Llera et al; IC6**

IC6 was reported by (Llera et al., 2019) and corresponds closely to IC81 from the current study (main analysis) and IC5 structural only analysis. A and D show uncorrected behavioural associations with IC81 and IC5, respectively. Tiles are coloured according to their correlation coefficient (r). * $p_{uncorrected} < 0.05$, ** $p_{uncorrected} < 0.01$, *** $p_{uncorrected} < 0.001$. B and C are scatter plots of the associations IC6 subject course and the subject courses of IC81 and IC5, respectively. Correlations (Spearman) were significant based on permutation testing ($p_{perm} < 0.001$). Calculations are based on n=207 individuals that overlapped between study cohorts.

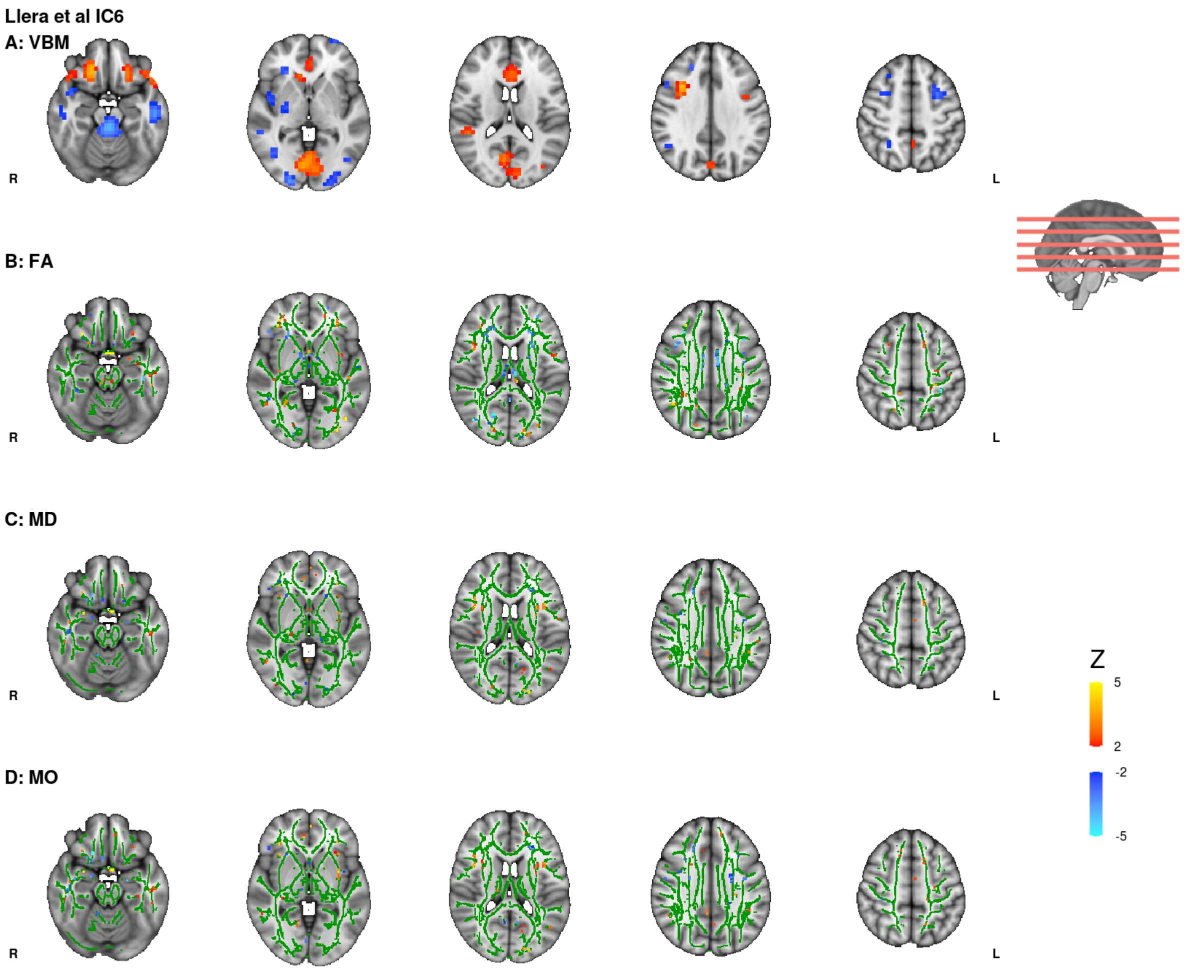


**Figure S16 spatial maps from Llera et al; IC6**

Spatial maps, for select modalities, generated in the analysis by (Llera et al., 2019) are presented for IC6. This is seen to correspond closely to IC81 (main analysis) and IC5 (structural analysis) from the current study. (A-D) relate to the VBM, FA, MD and MO modalities, respectively for Llera's IC6. Only VBM and DWI modalities are visualised. Colour gradient (blue-yellow) represents the Z score of the spatial map. Maps were thresholded at $|z|>2$. Sagittal brain slice shows location of axial slices used for each modality. Spatial maps are projected on the MNI T1 anatomical brain image for reference. DWI maps (FA, MD, MO) are displayed on top of the binarized FMRIB58 FA skeleton (green) as well as the MNI T1 template.

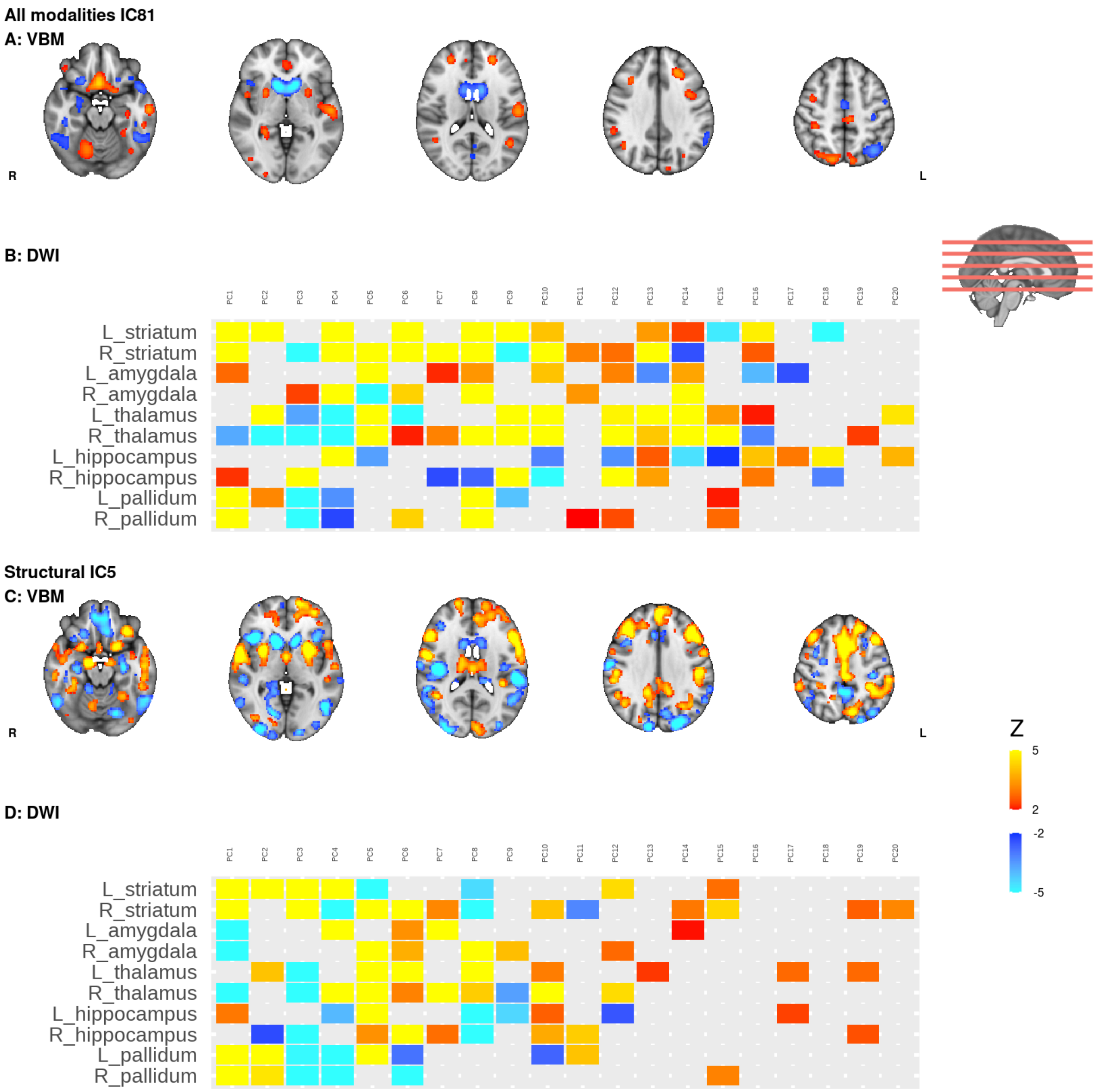


**Figure S17 spatial maps from IC81 (all modalities) and IC5 (structural)**

Spatial maps, for select modalities, generated in the current study are presented for IC81 (main analysis with all modalities) and IC5 (structural analysis). These are seen to correspond closely to IC6 from Llera et al. (A-B) relate to the VBM and DWI modalities, respectively for IC81. (C-D) similarly relate to the VBM and DWI modalities, respectively for IC5. Only VBM and DWI modalities are visualised. Colour gradient (blue-yellow) represents the Z score of the spatial map. Maps were thresholded at |z|>2. Sagittal brain slice shows location of axial slices used for VBM visualisation. VBM maps are projected on the MNI T1 anatomical brain image for reference. DWI maps display PCs 1-20.
